# R/qtl2: software for mapping quantitative trait loci with high-dimensional data and multi-parent populations

**DOI:** 10.1101/414748

**Authors:** Karl W. Broman, Daniel M. Gatti, Petr Simecek, Nicholas A. Furlotte, Pjotr Prins, Śaunak Sen, Brian S. Yandell, Gary A. Churchill

**Affiliations:** Department of Biostatistics and Medical Informatics, University of Wisconsin–Madison, Madison, Wisconsin 53706; Department of Horticulture, University of Wisconsin–Madison, Madison, Wisconsin 53706; Department of Statistics, University of Wisconsin–Madison, Madison, Wisconsin 53706; 23andMe, Mountain View, California 94043; The Jackson Laboratory, Bar Harbor, Maine 04609; Department of Genetics, Genomics, and Informatics, University of Tennessee Health Sciences Center, Memphis, Tennessee 38163; Department of Preventive Medicine, University of Tennessee Health Sciences Center, Memphis, Tennessee 38163; Center for Molecular Medicine, University Medical Center Utrecht, 3584CT Utrecht, The Netherlands

**Keywords:** software, QTL, multi-parent populations, MAGIC, Diversity Outbred mice, heterogeneous stock, Collaborative Cross

## Abstract

R/qtl2 is an interactive software environment for mapping quantitative trait loci (QTL) in experimental populations. The R/qtl2 software expands the scope of the widely-used R/qtl software package to include multi-parent populations derived from more than two founder strains, such as the Collaborative Cross and Diversity Outbred mice, heterogeneous stocks, and MAGIC plant populations. R/qtl2 is designed to handle modern high-density genotyping data and high-dimensional molecular phenotypes including gene expression and proteomics. R/qtl2 includes the ability to perform genome scans using a linear mixed model to account for population structure, and also includes features to impute SNPs based on founder strain genomes and to carry out association mapping. The R/qtl2 software provides all of the basic features needed for QTL mapping, including graphical displays and summary reports, and it can be extended through the creation of add-on packages. R/qtl2 comes with a test framework and is free and open source software written in the R and C++ programming languages.

## Introduction

There has been a resurgence of interest in the mapping of quantitative trait loci (QTL) in experimental organisms, spurred in part by the use of gene expression phenotypes (eQTL mapping; see Albert and Kruglyak 2015) to more rapidly identify the underlying genes, and by the development of multi-parent populations (de Koning and McIntyre 2017), including heterogeneous stocks (Mott *et al*. 2000; Mott and Flint 2002), MAGIC lines (Cavanagh *et al*. 2008; Kover *et al*. 2009), the Collaborative Cross (Churchill *et al*. 2004), and Diversity Outbred mice (Churchill *et al*. 2012; Svenson *et al*. 2012).

Multi-parent populations (MPPs) are genetically mixed populations derived from a small set of known founders that are typically but not necessarily inbred strains. The presence of multiple founder alleles imparts unique features to MPPs with significant advantages over traditional two-parent crosses. Allelic series of linked functional variants produce information-rich patterns of effects that can help identify causal variants and distinguish pleiotropy from chance co-localization of multiple QTL (King *et al*. 2012). MPPs provide high-resolution mapping, which results in fewer candidate genes and minimizes the confounding effects of linked loci. MPPs create new multi-locus allelic combinations by mixing founder genomes. The founder strain genomes of many MPPs have been or will be sequenced, and using high-density genotyping we can then accurately impute whole genomes of individuals (Oreper *et al*. 2017).

MPPs can be generated by many different breeding designs and have been developed in different model organisms including rats (Solberg Woods and Mott 2017), Drosophila (King *et al*. 2012), C. elegans (Noble *et al*. 2017), as well as a variety of plant species (Kover *et al*. 2009; DellAcqua *et al*. 2015; Bandillo *et al*. 2013; Huang *et al*. 2012a). Different breeding designs of MPPs give rise to different population structures and thus will require a flexible and general framework for analysis. The key challenges that arise in the analysis of MPP data include the reconstruction of the founder haplotype mosaic, imputation of whole-genome genetic variants, and analysis methods that can handle the multiple founder alleles and account for population structure.

There are numerous software packages for QTL mapping in classical two-parent experimental populations, including Mapmaker/QTL (Lincoln and Lander 1990), QTL Cartographer (Basten *et al*. 2002), R/qtl (Broman *et al*. 2003; Broman and Sen 2009), and MapQTL (Van Ooijen 2009). There are a smaller number of packages for QTL analysis in multi-parent populations, including DOQTL (Gatti *et al*. 2014), HAPPY (Mott *et al*. 2000), and mpMap (Huang and George 2011). Our aim in developing R/qtl2 is to provide an open-source, extensible software environment for QTL mapping and associated data analysis tasks that applies to the full range of classical and MPP cross designs.

The original R/qtl (hereafter, R/qtl1) is widely used, and has a number of advantages compared to proprietary alternatives. R/qtl1 includes a quite comprehensive set of QTL mapping methods, including multiple-QTL exploration and model selection (Arends *et al*. 2010; Broman and Speed 2002; Manichaikul *et al*. 2009), as well as extensive visualization and data diagnostics tools (Broman and Sen 2009). Further, users and developers both benefit by it being an add-on package for the general statistical software, R (R Core Team 2018). A number of other R packages have been written to work in concert with R/qtl1, including ASMap (Taylor and Butler 2017), ctl (Arends *et al*. 2016), dlmap (Huang *et al*. 2012b), qtlcharts (Broman 2015), vqtl (Corty and Valdar 2018), and wgaim (Taylor and Verbyla 2011).

R/qtl1 has a number of limitations (see Broman 2014), the most critical of which is that the central data structure generally limits its use to biparental crosses. Also, R/qtl1 was designed at a time when a dataset with more than 100 genetic markers was considered large.

Rather than extend R/qtl1 for multi-parent populations, we decided to start fresh. R/qtl2 is a completely redesigned R package for QTL analysis that can handle a variety of multi-parent populations and is suited for high-dimensional genotype and phenotype data. To handle population structure, QTL analysis may be performed with a linear mixed model that includes a residual polygenic effect. The R/qtl2 software is available from its web site (https://kbroman.org/qtl2) as well as GitHub (https://github.com/rqtl/qtl2).

## Features

QTL analysis in multi-parent populations can be split into two parts: calculation of genotype probabilities using multipoint SNP genotypes, and the genome scan to evaluate the association between genotype and phenotype, using those probabilities. We use a hidden Markov model (HMM; see Broman and Sen 2009, App. D) for the calculation of genotype probabilities. The HMM implemented in R/qtl2 is generalized from the implementation in R/qtl1 to accommodate the MPP founder haplotype structure. As the source of genotype information, R/qtl2 considers array-based SNP genotypes. At present, we focus solely on marker genotypes rather than array intensities, as in DOQTL, or allele counts/dosages from genotyping-by-sequencing (GBS) assays.

R/qtl2 includes implementations of many classical two-way crosses (backcross, intercross, doubled haploids, two-way recombinant inbred lines by selfing or sibling mating, and two-way advanced intercross populations), and many different types of multi-parent populations (4- and 8-way recombinant inbred lines by sibling mating; 4-, 8-, and 16-way recombinant inbred lines by selfing; 3-way advanced intercross populations, Diversity Outbred mice, heterogeneous stocks, 19-way MAGIC lines like the Kover *et al*. (2009) Arabidopsis lines, and 6-way doubled haploids following a design of maize MAGIC lines being developed at the University of Wisconsin–Madison).

A key component of the HMM is the transition matrix (or “step” probabilities), which are specific to the cross design. Transitions represent locations where the ancestry of chromosomal segments change from one founder strain haplotype to another. The transition probabilities for multi-way recombinant inbred lines are taken from Broman (2005). The transition probabilities for heterogeneous stocks and Diversity Outbred mice are taken from Broman (2012b), which uses the results of Broman (2012a).

The output of the HMM is a list of 3-dimensional arrays, one per chromosome, with dimensions corresponding to individuals x genotypes x marker loci. Array elements represent genotype probabilities that can reflect both the uncertainty of haplotype inference and the heterozygosity. The size and structure of the genotype dimension determine the form of the regression model that will be used in the genome scanning step. Thus once the genotype probabilities are defined, there is no need to reference the breeding scheme that gave rise to the cross population. For breeding schemes that are not currently implemented in the R/qtl2 HMM, the user can pre-compute and import a custom genotype probability data structure.

At present, R/qtl2 assumes dense marker information and a low level of uncertainty in the haplotype reconstructions, so that we may rely on Haley-Knott regression (Haley and Knott 1992) for genome scans to establish genotype-phenotype association. This may be performed either with a simple linear model (as in Haley and Knott 1992), or with a linear mixed model (Yu *et al*. 2006; Kang *et al*. 2008; Lippert *et al*. 2011) that includes a residual polygenic effect to account for population structure. The latter may also be performed using kinship matrices derived using the “leave-one-chromosome-out” (LOCO) method (see Yang *et al*. 2014).

To establish statistical significance of evidence for QTL, accounting for a genome scan, R/qtl2 facilitates the use of permutation tests (Churchill and Doerge 1994). For multi-parent populations with analysis via a linear mixed model, we permute the rows of the haplotype reconstructions as considered in Cheng and Palmer (2013). R packages such as qvalue (Storey *et al*. 2018) can be used to implement multiple-test corrections for high-dimensional data analysis (Storey 2002, 2003) such as gene expression QTL (eQTL) mapping.

R/qtl2 includes a variety of data diagnostic tools, which can be particularly helpful for data on multi-parent populations where the SNP genotypes are incompletely informative (i.e., SNP genotypes do not fully define the corresponding founder haplotype). These include SNP genotyping error LOD scores (Lincoln and Lander 1992) and estimated crossover counts.

## Examples

R/qtl2 reproduces the functionality of DOQTL (Gatti *et al*. 2014) but targets a broader set of multi-parent populations, in addition to Diversity Outbred mice. (DOQTL will ultimately be deprecated and replaced with R/qtl2.) Fig. 1 contains a reproduction, using R/qtl2, of Fig. 5 from Gatti *et al*. (2014). This is a QTL analysis of constitutive neutrophil counts in 742 Diversity Outbred mice (from generations 3–5) that were genotyped with the first generation Mouse Universal Genotyping Array (MUGA) (Morgan *et al*. 2016), which contained 7,851 markers, of which we are using 6,413.

**Figure 1:**
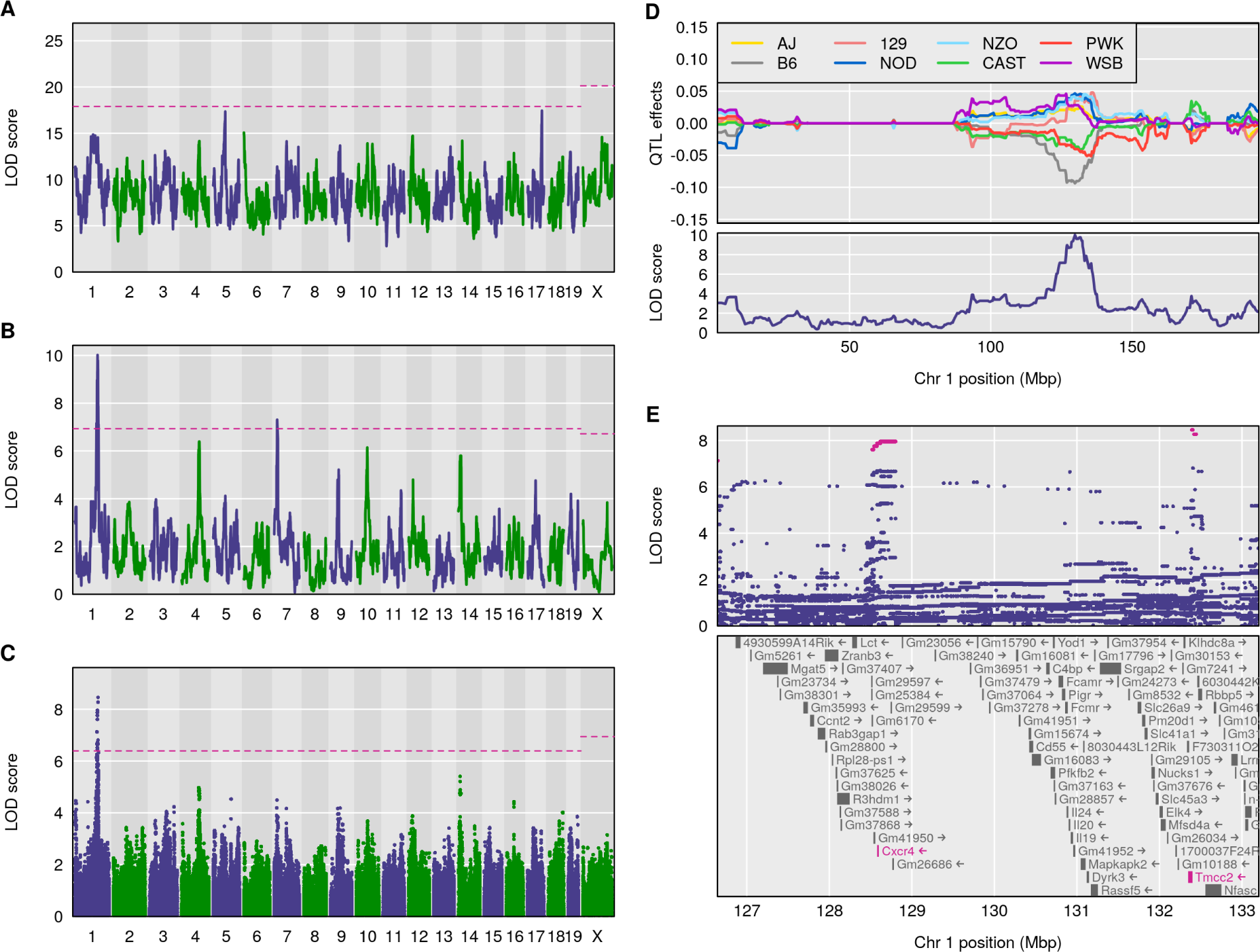
Reconstruction of Fig. 5 from Gatti *et al*. (2014), on the mapping of constitutive neu-trophil counts in 742 Diversity Outbred mice. (A) LOD scores from a genome scan using the full model (comparing all 36 genotypes for the autosomes and 44 genotypes for the X chromosome); the dashed horizontal line indicates the 5% genome-wide significance threshold, based on a per-mutation test. (B) LOD scores from a genome scan with an additive allele model (compare the 8 founder haplotypes). (C) LOD scores from a SNP association scan, using all SNPs that were genotyped in the eight founder lines. (D) Best linear unbiased predictors (BLUPs) of the eight haplotype effects in the additive model, along with the LOD curve on chromosome 1. (E) SNP association results in the region of the chromosome 1 QTL, along with genes in the region; SNPs with LOD scores within 1.5 of the maximum are highlighted in pink. All figures are produced with R/qtl2.

The regression model that R/qtl2 applies in a genome scan is determined by the HMM output in the genotype probabilities data structure. For an 8-parent MPP such as the DO mice, there are 36 possible diplotypes (44 on the X chromosome) and the genome scan will be based on a regression model with 35 degrees of freedom. With so many degrees of freedom, the model typically lacks power to detect QTL. An alternative representation collapses the 36 states to 8 founder “dosages” and uses a regression model with 7 degrees of freedom, assuming that the founder effects are additive at any given locus. R/qtl2 has the ability to incorporate SNP (and other variant) data from founder strains and to impute biallelic genotypes for every SNP. The genome scan on imputed SNPs is equivalent to an association mapping scan and can employ a additive (one degree of freedom) or general (two degrees of freedom) regression model.

Fig. 1A contains the LOD curves from a genome scan using a full model comparing all 36 possible genotypes with log neutrophil count as the phenotype and with sex and log white blood cell count as covariates. The horizontal dashed line indicates the 5% genome-wide significance level, derived from a permutation test, with separate thresholds for the autosomes and the X chromosome, using the technique of Broman *et al*. (2006). Fig. 1B contains the LOD curves from a genome scan using an additive allele model (corresponding to a test with 7 degrees of freedom), and Fig. 1C contains a SNP association scan, using a test with 2 degrees of freedom. All of these analyses use a linear mixed model with kinship matrices derived using the “leave-one-chromosome-out” (LOCO) method.

Fig. 1D shows the estimated QTL effects, assuming a single QTL with additive allele effects on chr 1, and sliding the position of the QTL across the chromosome. This is analogous to the estimated effects in Fig. 5D of Gatti *et al*. (2014), but here we present Best Linear Unbiased Predictors (BLUPs), taking the QTL effects to be random effects. This results in estimated effects that have been shrunk towards 0, which helps to clean up the figure and focus attention on the region of interest.

Fig. 1E shows individual SNP association results, for the 6 Mbp region on chr 1 that contains the QTL. As with the DOQTL software, we use all available SNPs for which genotype data are available in the 8 founder lines, and impute the SNP genotypes in the Diversity Outbred mice, using the individuals’ genotype probabilities along with the founder strains’ SNP genotypes.

Fig. 1 shows a number of differences from the results reported in Gatti *et al*. (2014), including that we see nearly-significant loci on chr 5 and 17 in the full model (Fig. 1A), and we see a second significant QTL on chr 7 with the additive allele model (Fig. 1B). Also, in Fig. 1E, we see associated SNPs not just at *∼*128.6 Mbp near the *Cxcr4* gene (as in Gatti *et al*. 2014), but also a group of associated SNPs at *∼* 132.4 Mbp, near *Tmcc2*. The differences between these results and those of Gatti *et al*. (2014) are due to differences in genotype probability calculations; R/qtl2 appears to be more tolerant of SNP genotyping errors (data not shown).

To further illustrate the broad applicability of R/qtl2, we reanalyzed the data of Gnan *et al*. (2014) on seed weight, seed number, and fruit length in 677 19-way Arabidopsis MAGIC lines from Kover *et al*. (2009). In Fig. 2, we show LOD scores for three traits and effect estimates for a selected QTL for each trait, as derived from the log P-values provided by Gnan *et al*. (2014) and as calculated with R/qtl2.

**Figure 2:**
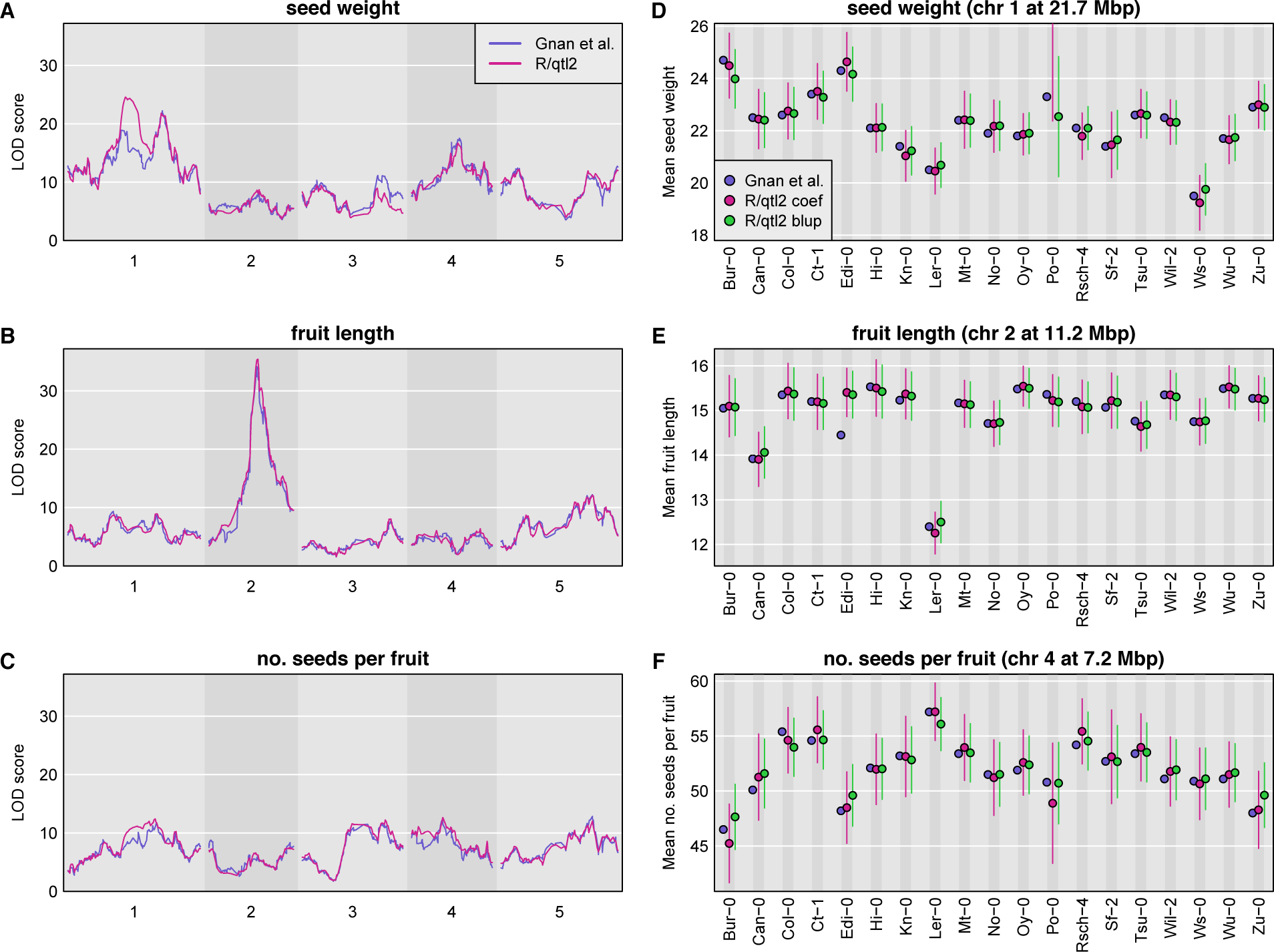
Analysis of 19-way Arabidopsis MAGIC data from Gnan *et al*. (2014) for three traits. The left panels show LOD curves with the results from Gnan *et al*. (2014) in blue, and from R/qtl2 in pink. The right panels show estimated QTL effects from Table 5 of Gnan *et al*. (2014) (blue), by linear regression from R/qtl2 (pink), and BLUPS from R/qtl2 (green).

The genome scan results are largely concordant except for an important difference in the LOD curve on chromosome 1 for seed weight (Fig. 2A). There are also smaller differences on chromosome 3 for seed weight (Fig. 2A) and chromosome 1 for number of seeds per fruit (Fig. 2C). These differences are likely due to differences in the calculated genotype probabilities, and deserve further study.

The estimated effects at the selected QTL are largely concordant (Fig. 2D–2F), but note that for the seed weight trait (Fig. 2D), R/qtl2’s estimate of the average seed weight for lines with the Po-0 allele is 39.9, well outside the plotted range. At this QTL, it appears that the 677 MAGIC lines all have small probabilities for carrying the Po-0 allele. The only other large difference is in Fig. 1E for fruit length, where the value reported in Gnan *et al*. (2014) for the Edi-0 allele is much smaller than that obtained with R/qtl2. Finally, note that throughout, the BLUPs are all shifted towards the mean, and that this shift is much larger for seed number (Fig. 1F) versus fruit length (Fig. 1E).

### Data and software availability

The data for Fig. 1 are available at the Mouse Phenotype Database (https://phenome.jax.org/projects/Gatti2). The data for Fig. 2 are available as supplemental files for Gnan *et al*. (2014) (https://doi.org/10.1534/genetics.114.170746). R/qtl2 input files for both datasets are available at GitHub (https://github.com/rqtl/qtl2data).

The R/qtl2 software is available from its web site (https://kbroman.org/qtl2) as well as GitHub (https://github.com/rqtl/qtl2). The software is licensed under the GNU General Public License version 3.0.

The code to create Fig. 1 and 2 is available at GitHub at https://github.com/kbroman/Paper_Rqtl2.

## Implementation

R/qtl2 is developed as a free and open source add-on package for the general statistical software, R (R Core Team 2018). Much of the code is written in R, but computationally intensive aspects are written in C++. (Computationally intensive aspects of R/qtl1 are in C.) We use Rcpp (Eddelbuettel and Francois 2011; Eddelbuettel 2013) for the interface between R and C++, to simplify code and reduce the need for copying data in memory. We use roxygen2 (Wickham *et al*. 2017) to develop the R package documentation.

Linear algebra calculations, such as matrix decomposition and linear regression, are a central part of QTL analysis. We use RcppEigen (Bates and Eddelbuettel 2013) and the Eigen C++ library (Guennebaud *et al*. 2010) for these calculations. For the fit of linear mixed models, to account for population structure with a residual polygenic effect, we closely followed code from PyLMM (Furlotte 2015). In particular, we use the basic technique described in Kang *et al*. (2008), of taking the eigen decomposition of the kinship matrix.

In contrast to R/qtl1, which includes almost no formal software tests, R/qtl2 includes extensive unit tests to ensure correctness. We use the R package testthat (Wickham 2011) for this purpose. The use of unit tests helps us to catch bugs earlier, and revealed several bugs in R/qtl1.

## Discussion

We have completed the core of the R/qtl2 software package, which is a re-implementation of the widely-used software R/qtl, to better handle high-dimensional genotypes and phenotypes, and modern cross designs including MPPs. This software forms a key computational platform for QTL analysis in MPPs, and includes genotype reconstruction for a variety of MPP designs (including MAGIC lines, the Collaborative Cross, Diversity Outbreds, and heterogeneous stock), numerous facilities for quality-control assessments, QTL genome scans by Haley-Knott regression (Haley and Knott, 1992) and linear mixed models to account for population structure, and BLUP-based estimates of QTL effects. Most procedures in R/qtl2 can make use of the multiple CPU cores on a given machine, to speed computations by parallel processing.

While the basic functionality of R/qtl2 is complete, there are a number of areas for further development. In particular, we would like to further expand the set of crosses that may be considered, including partially-inbred recombinant inbred lines (so that we may deal with residual heterozygosity, which presently is ignored). We have currently been focusing on exact calculations for specific designs, but the mathematics involved can be tedious. We would like to have a more general approach for genotype reconstruction in multi-parent populations, along the lines of RABBIT (Zheng *et al*. 2015) or STITCH (Davies *et al*. 2016). Plant researchers have been particularly creative in developing unusual sets of MAGIC populations, and by our current approach, each design requires the development of design-specific code, which is difficult to sustain. In addition, we will provide facilities for importing data in more general formats, including genotype probabilities/reconstructions and kinship matrices that were derived from other software packages. This will further expand the scope for R/qtl2 by making its QTL analysis facilities usable beyond the set of MPP designs that can be handled by our genotype reconstruction code.

Another important area of development is the handling of genotyping-by-sequencing (GBS) data. We are currently focusing solely on called genotypes. With low-coverage GBS data, it is difficult to get quality genotype calls at individual SNPs, and there will be considerable advantage to using the pairs of allele counts and combining information across SNPs. Extending the current HMM implementation in R/qtl2 to handle pairs of allele counts for GBS data appears straightforward.

At present, QTL analysis in R/qtl2 is solely by genome scans with single-QTL models. Consideration of multiple-QTL models will be particularly important for exploring the possibility of multiple causal SNPs in a QTL region, along the lines of the CAVIAR software (Hormozdiari *et al*. 2014).

We have currently focused solely on Haley-Knott regression (Haley and Knott 1992) for QTL analysis. This has a big advantage in terms of computational speed, but it does not fully account for the uncertainty in genotype reconstructions. But the QTL analysis literature has a long history of methods for dealing with this genotype uncertainty, including interval mapping (Lander and Botstein 1989) and multiple imputation (Sen and Churchill 2001). While this has not been a high priority in the development of R/qtl2, ultimately we will include implementations of these sorts of approaches, to better handle regions with reduced genotype information.

We will continue to focus on lean implementations of fitting algorithms, such as simple linear mixed models with a single random effect for kinship, that will be widely used for genome-wide scans. But we will also seek to simplify the use of external packages, for genome scans with more complex models.

R/qtl2 is an important update to the popular R/qtl software, expanding the scope to include multi-parent populations, providing improved handling of high-dimensional data, and enabling genome scans with a linear mixed model to account for population structure. R/qtl1 served as an important hub upon which other developers could build; we hope that R/qtl2 can serve a similar role for the genetic analysis of multi-parent populations.

## Acknowledgments

This work was supported in part by National Institutes of Health grants R01GM074244 (to K.W.B.), R01GM070683 (to K.W.B. and G.A.C.), and R01GM123489 (to S.S.). The authors thank Paula Kover for assistance with the data from Gnan *et al*. (2014).

